# Concurrent EEG- and fMRI-derived functional connectomes exhibit linked dynamics

**DOI:** 10.1101/464438

**Authors:** Jonathan Wirsich, Anne-Lise Giraud, Sepideh Sadaghiani

## Abstract

Connectivity across distributed brain regions commonly measured with functional Magnetic Resonance Imaging (fMRI) exhibits infraslow (<0.1Hz) spatial reconfigurations of potentially critical importance to cognition. Cognitively relevant neural communication, however, employs synchrony at fast speeds. It is unclear how fast oscillation-coupling across the whole-brain connectome relates to connectivity changes in fMRI, an indirect measure of neural activity. In two datasets, electroencephalography (EEG) revealed that synchronization in all canonical oscillation-bands reconfigures at infraslow speeds, coinciding with connectivity changes in concurrently recorded fMRI in corresponding region-pairs. The cross-modal tie of connectivity dynamics was widely distributed across the connectome irrespective of EEG frequency-band. However, the cross-modal tie was strongest in visual to somatomotor connections for slower EEG-bands, and in connections involving the Default Mode Network for faster EEG-bands. The findings provide evidence that functionally relevant neural synchrony in all oscillation-bands slowly reconfigures across the whole-brain connectome, and that fMRI can reliably measure such dynamics.

## Introduction

To date, our knowledge about the topography of neural communication in the human brain is largely derived from fMRI, an indirect measure of neural activity. Using fMRI, the brain has been characterized in terms of different large-scale intrinsic connectivity networks (ICNs) (1–3). Collectively, the connectivity within and between ICNs can be represented as a whole-brain connectivity graph, or connectome (4). Extensive efforts have been undertaken towards establishing the neural origin of this functional connectivity (FC) organization (5–11). This line of research has traditionally focused on static properties of FC by averaging brain activity across the entire recording period. To date, a comparable spatial organization of static whole-brain FC has been observed across EEG and fMRI (6, 11), MEG and fMRI (5, 8) and intracranial EEG and fMRI (7, 9). Because cognitive processes are inherently dynamic, time-varying reconfigurations across the fMRI-derived functional connectome occurring at infraslow speeds (typically defined as < 0.1Hz (12, 13)) are receiving increasing attention (14, 15). Such reconfigurations or dynamics of functional connectivity (dFC) are suggested to represent flexible changes across distinct cognitive architectures that support different cognitive processes (16). This intriguing possibility derived from fMRI could explain the role of intrinsic connectivity in cognition (17, 18). However, the relationship between fMRI-derived dFC and cognitively relevant electrophysiological mechanisms of information exchange, specifically oscillatory synchrony (19, 20), has not yet been substantiated in direct measures of neural connectivity across the connectome.

Such direct neural measures are essential because investigations of time-varying fMRI connectivity have been plagued by concerns about physiological noise, head motion, and sampling variability (21, 22), as well as uncertainty regarding how to adequately model and account for such spurious ‘dynamics’ (15, 23). In particular, three questions remain unanswered in direct neurophysiological recordings. Does neurophysiological connectivity (phase coupling) typically derived from fast neural oscillations (~1-100 Hz) fluctuate in strength at infraslow speeds (< 0.1Hz) across regions of the connectome? If so, do such infraslow changes in neurophysiological connectivity co-occur with fMRI-derived connectivity changes across corresponding connectome regions? And to what extent is the putative cross-modal link frequency-dependent? More specifically, changes in fMRI-derived connectivity between a given set of brain regions could be accompanied by infraslow connectivity changes in all canonical oscillation frequencies. Alternatively, fMRI connectivity dynamics could be accompanied by fluctuations of different frequency bands across different sets of brain regions. In the following sections, we discuss the status quo and knowledge gap regarding these questions. Since studying the temporal relationship of dFC between fMRI and neurophysiological signals necessitates concurrently recorded data, the following overview focuses largely on multimodal research.

The dynamic modulation in phase coupling of neurophysiological oscillations as measured by EEG and MEG has been well-established as a mechanism for long-range neural communication (19, 20). However, it is unknown whether these neurophysiological connectivity dynamics extend to infraslow time scales comparable to those driving dFC in fMRI. Indications that this may be the case come from concurrent EEG-fMRI studies assessing neurophysiological phase coupling at sensor level and building a coarse global connectivity average across all EEG electrode pairs (24, 25). Such spatially averaged neurophysiological connectivity timecourses indeed exhibit fluctuations at infraslow timescales. However, connectivity across electrodes does not adequately reflect neurophysiological connectivity across brain regions (26), and global averaging further removes any spatial information. Additionally, these studies relate neurophysiological connectivity to fMRI amplitude but not fMRI connectivity.

Conversely, other concurrent EEG-fMRI studies have compared neurophysiological signal amplitude to fMRI-derived connectivity dynamics. EEG band-limited global field power (average across all electrodes) has been linked to fMRI-derived dFC both in specific ICNs (27) and the whole brain (28, 29). Similarly, Allen et al. (30) demonstrated that the reoccurring states of fMRI-based dFC co-occur with changes in the EEG power spectrum of certain electrodes. Again, no information about neurophysiological connectivity across brain regions can be obtained from these studies since they use sensor-level EEG amplitude. Thus, it remains unknown whether neurophysiological dFC across regions of the connectome changes in unison with the distributed dynamics observed in concurrent fMRI.

Another set of studies made the crucial advance to comparing fMRI connectivity with spatially localized neurophysiological connectivity obtained from invasive recordings. Neurophysiological oscillations recorded invasively in the rat show interhemispheric connectivity across homologous somatosensory areas that co-fluctuate with concurrent fMRI-derived dFC across the same regions (31, 32). Concurrent intracranial-EEG and fMRI in presurgical patients has revealed that region pairs showing high levels of connectivity dynamics in fMRI do so in invasive neurophysiological recordings as well (33). Unfortunately, the spatial coverage of such invasive electrophysiology studies is inherently limited. Consequently, they don’t inform about the correspondence of dynamic changes in the connectome’s whole-brain FC topography across neurophysiology and fMRI.

Closing this gap requires the study of neurophysiological connectivity dynamics across all regions of the whole-brain connectome concurrently with fMRI. Neurophysiological recordings over the scalp, i.e. EEG and MEG, are subject to limited spatial localizability. However, recent methodological advances have made it possible to measure whole-brain connectivity patterns and their dynamics from source-reconstructed signals, at least for coarse whole-brain parcellations (34, 35). The reliability of MEG- and EEG-derived source-localized connectomes is confirmed by significant spatial correspondence of their static connectivity patterns to both structural and fMRI-derived static connectomes (6, 11, 36, 37). However, no studies have investigated this whole-brain cross-modal correspondence in a dynamic framework, leaving the question of the relation between neurophysiological and fMRI-derived dFC across the connectome unanswered.

An interesting observation of the above-mentioned static cross-modal studies is that a stable connectome architecture comparable to that of fMRI exists for all canonical oscillation frequencies in EEG, albeit to varying degrees. Only two studies have compared fMRI-derived and concurrently recorded neurophysiological whole-brain connectomes (source-localized EEG). Deligianni et al. (6) found significant spatial similarity across whole-brain fMRI-derived FC and EEG-based band-limited amplitude-coupling, and this similarity was stronger for lower EEG frequencies than for β and γ bands. Investigating EEG phase-coupling rather than amplitude-coupling (11), we likewise observed significant spatial similarity between concurrent static EEG and fMRI connectomes for all oscillation bands, albeit weakening in the γ band (r>0.3 for δ through β bands, r=0.16 for γ). While the EEG frequency band with strongest static connectivity differed across connections (also cf.(8)), the connectivity pattern for a given band was not restricted to specific ICNs. Here, we seek to answer whether the putative time-varying co-evolution of fMRI and EEG dFC is likewise spatially extensive in all oscillatory frequency bands.

We hypothesize that dFC derived from source-space EEG shows infraslow changes that covary over time with fMRI dFC on a connection-wise basis across the whole brain. Further, we aim to characterize the whole-brain topographical organization of this cross-modal relationship for each canonical EEG frequency band.

## Results

We addressed the neurophysiological basis of fMRI dFC, its spatial topography, and its frequency-specificity in a primary concurrent EEG-fMRI dataset during task-free resting state (n=26, 3×10min), and assessed generalization in a second dataset (n=16, 10min). While eyes were closed in the first dataset, the second dataset was recorded with eyes open, allowing us to further assess generalization or differences across these conditions. After conservatively eliminating head motion based on both data modalities, preprocessed fMRI signal was averaged to the 68 cortical regions of the Desikan atlas (38, 39). Preprocessed EEG signals were source-reconstructed (40–42) to the same atlas (Fig. 1a).

### Static connectivity relationship across EEG and fMRI

First, we sought to confirm that the previously reported static relation between fMRI-derived and EEG-derived connection-wise connectivity strength (6, 11) holds true for the two datasets of this study. To this end, we assessed connectivity averaged across the total duration of EEG and fMRI data. Connectivity in source-projected EEG was quantified as band-limited (canonical frequency bands: δ, θ, α, β, γ) phase coupling using imaginary coherence (43, 44), and connectivity in fMRI was quantified as Pearson’s r for the same whole-brain parcellation atlas. In line with our prior work (11), static FC of all EEG bands were spatially correlated to FC of fMRI (main dataset fMRI vs. δ/θ/α/β/γ: r=0.34/0.34/0.33/0.36/0.29; generalization dataset fMRI vs. δ/θ/α/β/γ: r=0.34/0.33/0.36/0.41/0.39, taking the average FC for each connection across all subjects. See SI Fig. 1). This observation reaffirms the link between the spatial organization of connectivity across modalities, and confirms sufficient quality of EEG source localization to the whole-brain parcellation.

**Fig. 1:**
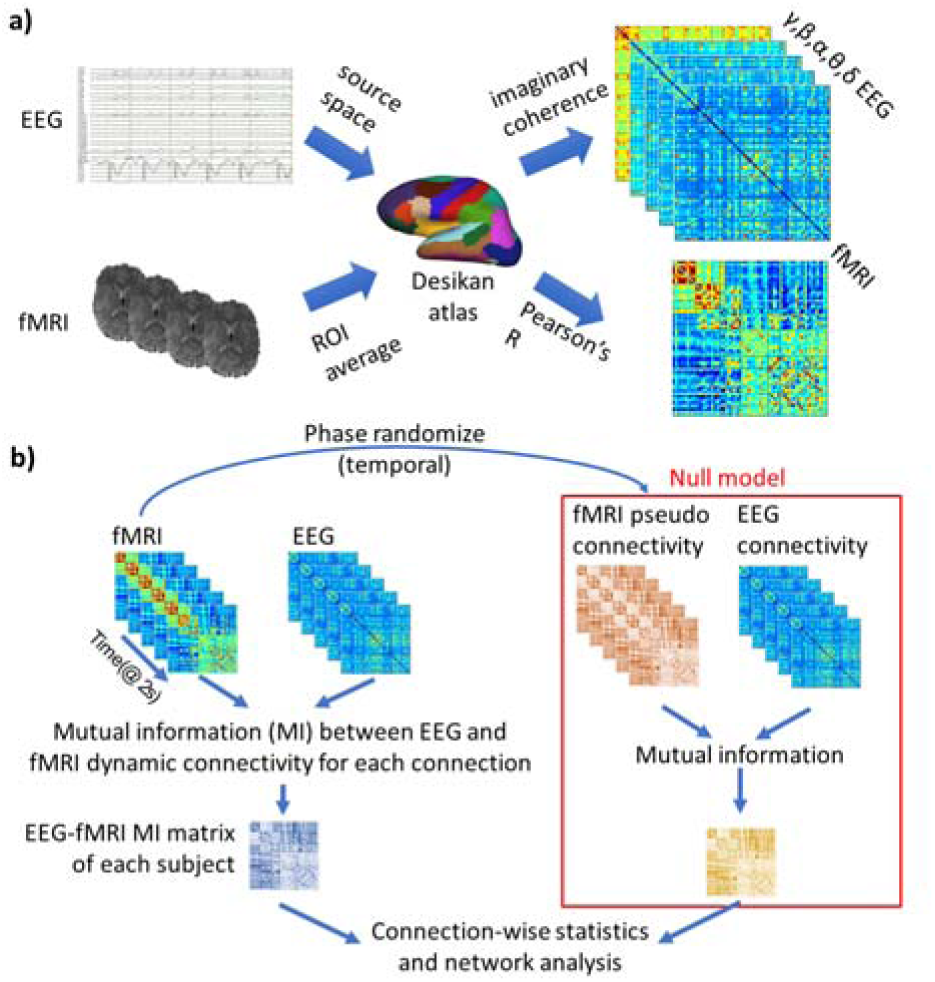
a) Construction of EEG and fMRI connectomes. EEG was source reconstructed and fMRI signal was averaged for the 68 cortical regions of the Desikan atlas (38). Pearson’s correlation of fMRI timecourses and imaginary coherence of band-limited EEG source signal for each region pair were used to build connectomes. b) In each data modality, dynamic FC was derived from a 1min window sliding at 2s (= repetition time of fMRI). For fMRI Pearson’s correlation at each connection was calculated over all samples of the 1-min window. For EEG, imaginary coherence was calculated at each connection in 2s segments (corresponding to the repetition time of fMRI), then averaged over the 1min window. To quantify the similarity of the connectivity dynamics across the data modalities, mutual information between the fMRI connectivity timecourse and the EEG connectivity timecourse was calculated at each connection. The ensuing mutual information at each connection was statistically compared to a null model. The null model consisted of the mutual information between the EEG connectivity timecourse and temporally phase scrambled fMRI connectivity timecourse (45 46)

### Dynamic connectivity relationship across EEG and fMRI

Next, we used a sliding window approach to assess whether time-varying changes in the whole-brain connectivity pattern derived from fMRI are linked to dynamics in band-limited phase coupling across modalities (Fig. 1b). We applied a 1min-wide window sliding at the resolution of the fMRI recordings in order to focus on infraslow dynamics characteristic of fMRI dFC (within the limits of methodologically reliable temporal resolution, as detailed in Methods). We used the information-theoretic measure of mutual information, which assesses the relationship between the modalities without assuming linearity. We tested these co-dynamics against a null model that randomizes the temporal phase of the fMRI connectivity timecourse for all connections (45, 46). SI Fig. 2 shows the distribution of mutual information of one randomly selected subject per dataset. Dynamic FC in δ, θ, α, β and γEEG showed high mutual information with fMRI connectome dynamics, significantly outperforming the null model in virtually all region-pairs: in 99.21% of connections for δEEG-fMRI, 98.82% in θEEG-fMRI, 98.2% in αEEG-fRMI, 96.88% in βEEG-fMRI and 98.24% in γEEG-fMRI (paired t-test, p<0.05, Bonferroni corrected for 2248 connections). This strong cross-modal relationship was confirmed in the generalization dataset, although it was significant in a smaller number of connections, especially for βEEG-fMRI and γEEG-fMRI: δEEG-fMRI=39.07%, θEEG-fMRI=30.64% αEEG-fRMI=27.22%, βEEG-fMRI=15.63%, γEEG-fMRI=13.04% (paired t-test, p<0.05, Bonferroni corrected for 2248 connections). The relative reduction in effect size in the replication dataset is in line with the smaller sample size and shorter recording duration. Fig. 2 shows the statistical outcome against the null model at a connection-wise resolution.

**Fig. 2:**
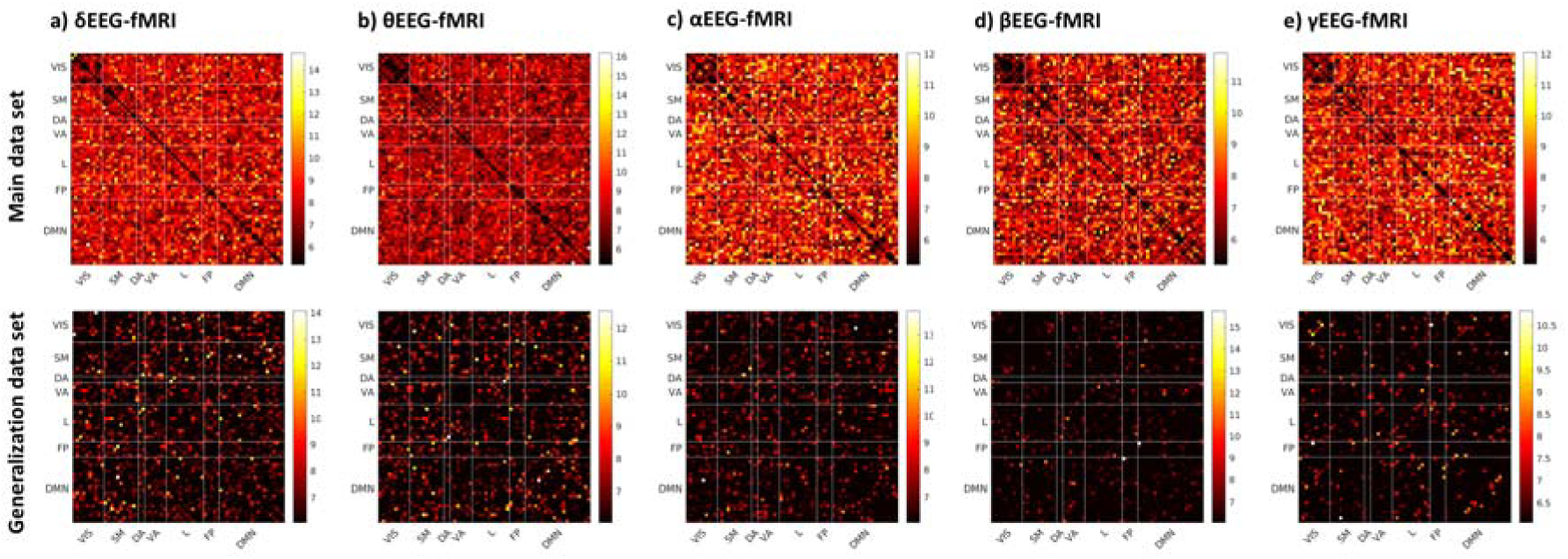
Distribution of t-values of the paired t-test mutual information across all connections between fMRI and EEG dFC time courses using real vs. phase-scrambled fMRI connectivity timecourses (null model) for each EEG band. Minimum t-values is defined by the Bonferroni corrected threshold t_25_=5.21 (main data set) and t_15_=6.06 (generalization data set). Almost all connections passed this threshold in the main dataset (>96.9% for all EEG bands). In the less extensive generalization dataset, a large proportion of connections passed the significance threshold, but the proportion progressively decreasing with increasing EEG frequency (39% to 13% from δ to γ). Connections are arranged according to canonical intrinsic networks (Visual (VIS), Somatomotor (SM), Dorsal Attention (DA), Ventral Attention (VA), Limbic (L), Fronto-Parietal (FP), and Default Mode Network (DMN)).

Additionally, we tested for generalizability at a connection-wise level. The connection-wise strength of EEG-fMRI mutual information averaged across all subjects was strongly correlated between primary and replication datasets for fMRI compared to δ/θ/αEEG (r=0.49/0.47/0.33), although very modest to no correlation was observed for fMRI vs. β/γEEG (r=0.06/-0.01, Fig. 3). A split-half approach indicated that the lack of connection-wise generalization for β and γ bands is due to the lower signal-to-noise ratio compared to slower frequencies especially in the less extensive generalization dataset (SI Results and Table S1).

**Fig. 3:**
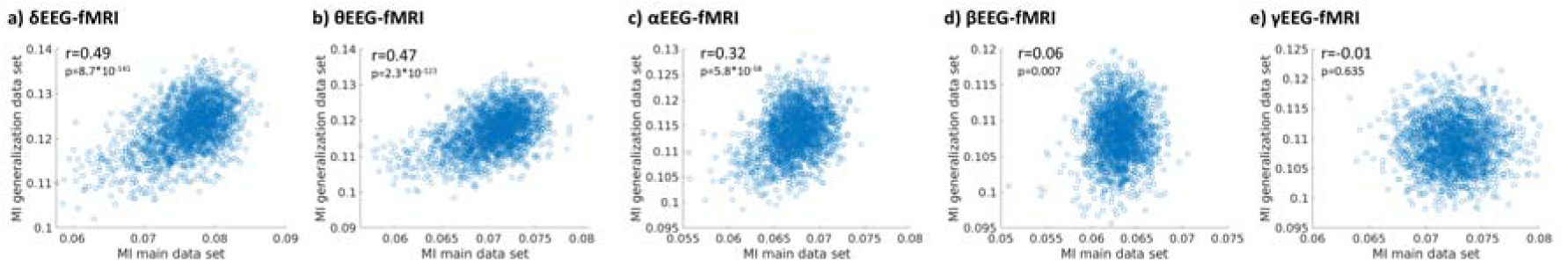
Connection-wise comparison of the EEG-fMRI relationship across the two datasets. Each data point reflects the mutual information between EEG dFC and fMRI dFC for a given connection averaged across all subjects of the respective datasets. Scatter plots are provided for the relation of fMRI dFC to a) δEEG, b) θEEG, c) αEEG, d) βEEG, and e) γEEG-dFC. The whole-brain distribution of mutual information correlated across the two datasets for δ, θ and αEEG.

In summary, fMRI signal correlations were linked to slow modulations of oscillatory phase-coupling across vast proportions of the connectome’s region-pairs. This was true for all canonical EEG frequency bands in both datasets. The strength of the EEG-fMRI relationship was correlated on a connection-wise basis between primary and generalization datasets in the δ, θ, and α bands.

### Spatial topography of the dynamic relationship

Next, we sought to characterize the spatial topography of co-dynamics beyond the above-described all-encompassing relation between EEG and fMRI dFC. Specifically, we assessed how the connections for which fMRI dFC was most strongly linked to EEG dFC were distributed over canonical ICNs (3) for each EEG frequency band. To this end, we selected the 200 connections with the strongest similarity of EEG and fMRI dFC over time, i.e. highest cross-modal mutual information (compared to the null data as assessed by the above-described t-statistic).

Fig. 4 visualizes the networks of the top-200 connections linked between fMRI dFC and EEG dFC for each canonical oscillation band. To understand the distribution of the top-200 connections with respect to ICNs, we mapped the connections to an atlas of seven canonical networks (Visual, Somatomotor, Dorsal Attention, Ventral Attention, Limbic, Fronto-Parietal, and Default Mode (DMN) networks (3), Fig 4a). The number of connections between any given pair of canonical ICNs is visualized in Fig 4b. This distribution of pairwise ICN connection density was strongly correlated across primary and generalization datasets for all bands (δ/θ/α/β/γ: r=0.72/0.84/0.80/0.87/0.71). We established that this distribution of the top-200 connections was not driven by the number of ICN nodes or other potential biases. For each ICN pair (e.g. DMN-Visual), we tested whether the number of connections was significantly higher than chance by randomly selecting (n=100,000) 200 connections to derive a statistic of the number of connections randomly mapping to each ICN-pair (Tables S3-S7). Interestingly, the dynamic reconfigurations of the top-200 connections with strongest tie across EEG and fMRI were situated between rather than within ICNs (off-diagonal compared to diagonal of matrices in 4b). A notable exception to this observation was the within-DMN connectivity, most notably in the β band. A high number (significantly above permutation chance) of connections with strongest cross-modal relationship connected DMN regions among themselves and to other ICNs, especially in α, β and γ frequencies. Further, the δ, θ and α frequencies showed strongest cross-modal link of FC dynamics between the Visual and Somatomotor networks. Additionally, a significantly high number of strong connections was observed beyond the above-described DMN-dependent connectivity and Visual-Somatomotor connectivity, most notably for Visual to Ventral Attention in δ, and Somatomotor to Dorsal Attention in θ and γ bands (Fig. 4b, Tables S3-S8).

When repeating the above-described permutation test in the generalization dataset (comparison of top-200 to 200 randomly selected connections), we replicated the large number of connections for Visual-Somatomotor in δ and θ bands, and for DMN-DMN in the β band (Tables S3-S7, green cells). The Somatomotor-Visual dominance did not replicate for the α band, presumably because of the unique sensitivity of this band to eyes closed vs. open conditions (main vs. generalization dataset, respectively). We performed an additional analysis to further ascertain that the topographic similarity of the top-200 connections across datasets was not due to chance. This permutation analysis (detailed in supplementary materials) found that the connection counts in the two datasets were drawn from the same distribution in all ICN pairs. Overall, these results support generalization of the spatial topography of co-dynamics between fMRI and EEG dFC at the coarser ICN-wise resolution.

**Fig. 4:**
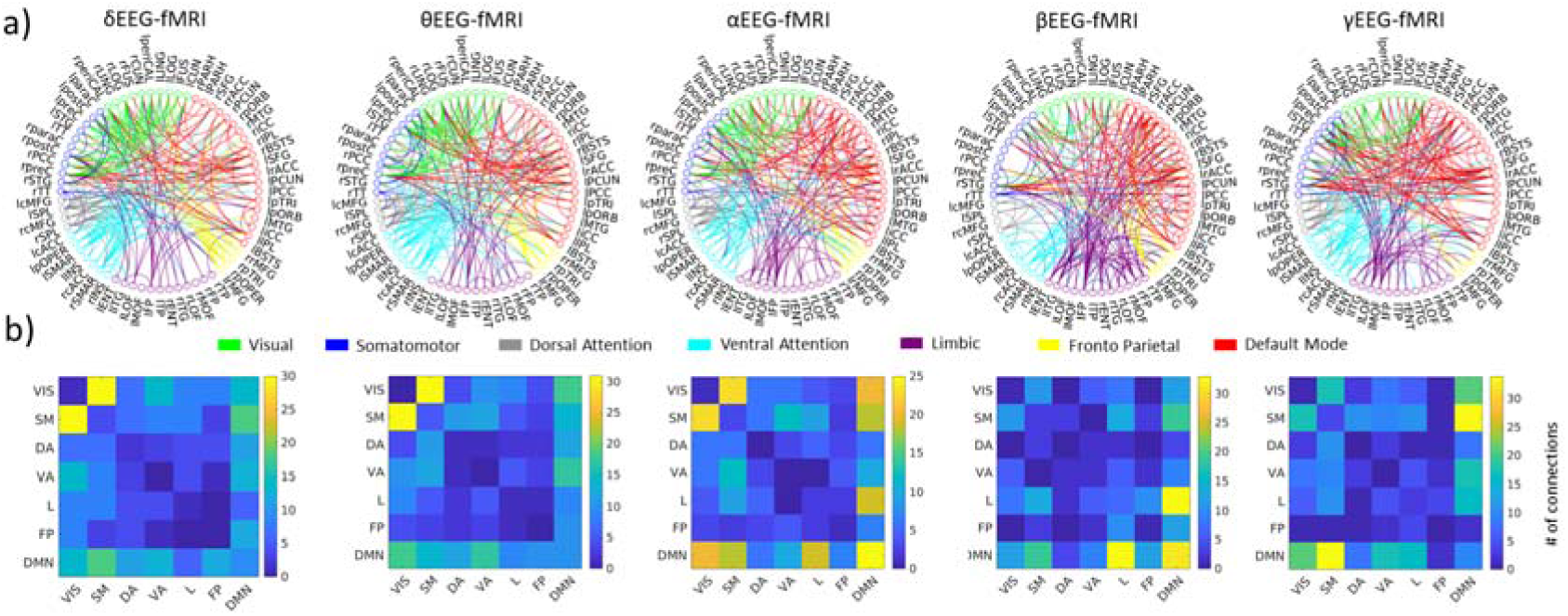
Spatial characterization of temporally linked EEG-fMRI connectivity dynamics. a) Topographical distribution of the 200 connections with the strongest EEG-fMRI dFC relationship for each EEG frequency band. Connections are color-coded according to the seven major canonical intrinsic networks (3). Individual region labels are listed in table S9. b) Mapping of those 200 connections to the intrinsic networks. The color scale depicts the count of connections falling between a given pair of canonical intrinsic networks (Visual (VIS), Somatomotor (SM), Dorsal Attention (DA), Ventral Attention (VA), Limbic (L), Frontoparietal (FP), and Default Mode Network (DMN)). The top-200 connections are dominated by within- and between-network connections of the DMN, especially for α, β and γ frequencies. Additionally, a large number of the top-200 connections fall between VIS and SM networks in the δ, θ and α frequencies. Results are visualized for the primary dataset, while a strongly correlated distribution over intrinsic networks was observed in the generalization dataset (Tables S3-S8).

In summary, while our first analysis showed a significant relationship between fMRI and EEG for a large proportion of connections and irrespective of oscillation frequency, the strength of this relationship varied over space depending on the EEG frequency. The connections with the strongest cross-modal relationship (top 200) for each EEG frequency band were positioned between a different set of ICNs. In particular, we observed a dominant role of the DMN -- especially in the faster frequency bands (β and γ) -- as well as a dominance of connections between the Visual and Somatomotor networks in slower frequency bands (δ and θ). The α band exhibited characteristics of both the faster and slower frequencies.

### Frequency-specificity of the dynamic relationship

Finally, we sought to directly and statistically corroborate the frequency-specificity of the cross-modal relationship on a connection-wise basis. To this end, we combined the EEG-fMRI mutual information matrices for all EEG bands into an ANOVA (5 levels for 5 frequency bands). First, we tested if the band-specific mutual information strength is driven by global shifts in the EEG connectivity due to head movement. To this end, we assessed the correlation of the fMRI-derived framewise displacement with the global EEG connectivity (average across connections) of each band in each run. We found no notable relationship between movement and EEG connectivity (all R<0.05; see SI: table S2). To further exclude global shifts that may be induced by parameters that are difficult to control (e.g. impact of the cardiobalistic artefact on the EEG connectome, and impact of the residual gradient artefact) and that may differentially affect each EEG band, we z-transformed each band’s mutual information matrix. As such, the connection-wise ANOVA assessed whether the connections with strongest mutual information in individual EEG bands overlap across bands, or instead are unique to specific bands. Statistical testing indicated a main effect of EEG band (Primary dataset: auxiliary uncorrected threshold F_4,125_>2.29; p<0.05, and p<0.05 NBS-corrected, with the significant cluster comprising 21.8% of connections; Generalization dataset: auxiliary uncorrected threshold F_4,75_>2.34; p<0.05, and p<0.05 NBS-corrected, comprising 14.9% of connections).

Exploratory post-hoc t-tests revealed that δ, β and γ band dFC are each organized in a frequency specific network with higher correspondence to fMRI dFC relative to all other EEG bands. Table 1 shows connection-wise t-thresholds resulting in a connected set of 100 region pairs at corrected statistical level (p<0.05, NBS-corrected). Fig. 5a visualizes this frequency-specific network for δ, β and γ bands in physical brain space, and Fig. 5b maps the connections according to canonical ICNs. Confirming the observations described above (Fig. 4b), networks of frequency-specific cross-modal dynamics consisted predominantly of DMN connections to itself and the rest of the brain. Additionally, the δ-specific set of connections comprised a high number of Somatomotor-Visual and Somatomotor-Frontoparietal connections, whereas the β-specific set showed a dominance of DMN-Limbic connections. Finally, the γ-specific set comprised a high number of DMN-Somatomotor and DMN-Limbic connections (Fig. 5c). Exploratory post-hoc analysis in the generalization dataset replicated a set of connections with significantly higher mutual information for δEEG-fMRI, βEEG-fMRI, and γEEG-fMRI (Table 1). We do not further discuss connections showing stronger cross-modal relationship in θ compared to other bands, since this result did not replicate in the generalization dataset. SI Fig. 3 demonstrates that the top-100 connection with strongest tie between fMRI dFC and δ-, β- and γ-band dFC are largely non-overlapping in both main and generalization datasets. To conclude, when contrasted directly across frequency bands, frequency-specificity of the EEG-fMRI relationship was confirmed with a DMN (especially to Limbic) and Somatomotor-Visual dominance.

**Table 1:**
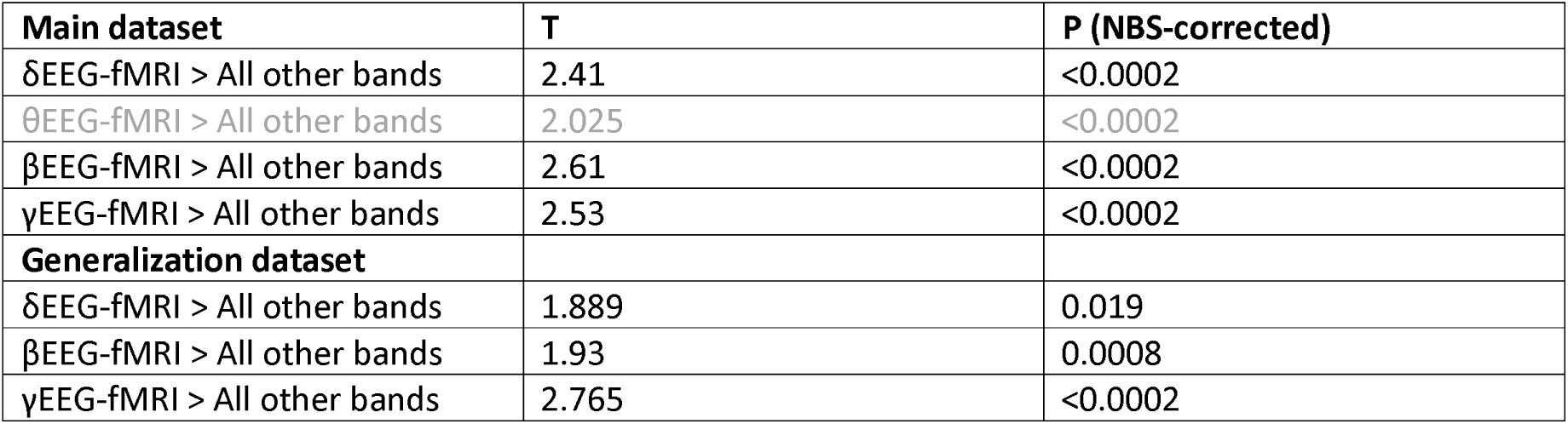
NBS (5000 iterations) shows a network of significantly increased mutual information for δEEG-fMRI and γEEG-fMRI (connection-wise T-threshold is chosen to limit the network size to 100 connections, one sided t-test one band to all the others). The ensuing top 100 connections are visualized in Fig. 5.

| Main dataset | T | P (NBS-corrected) |
| --- | --- | --- |
| δEEG-fMRI > All other bands | 2.41 | <0.0002 |
| θEEG-fMRI > All other bands | 2.025 | <0.0002 |
| βEEG-fMRI > All other bands | 2.61 | <0.0002 |
| γEEG-fMRI > All other bands | 2.53 | <0.0002 |
| **Generalization dataset** |  |  |
| δEEG-fMRI > All other bands | 1.889 | 0.019 |
| βEEG-fMRI > All other bands | 1.93 | 0.0008 |
| γEEG-fMRI > All other bands | 2.765 | <0.0002 |

**Fig. 5:**
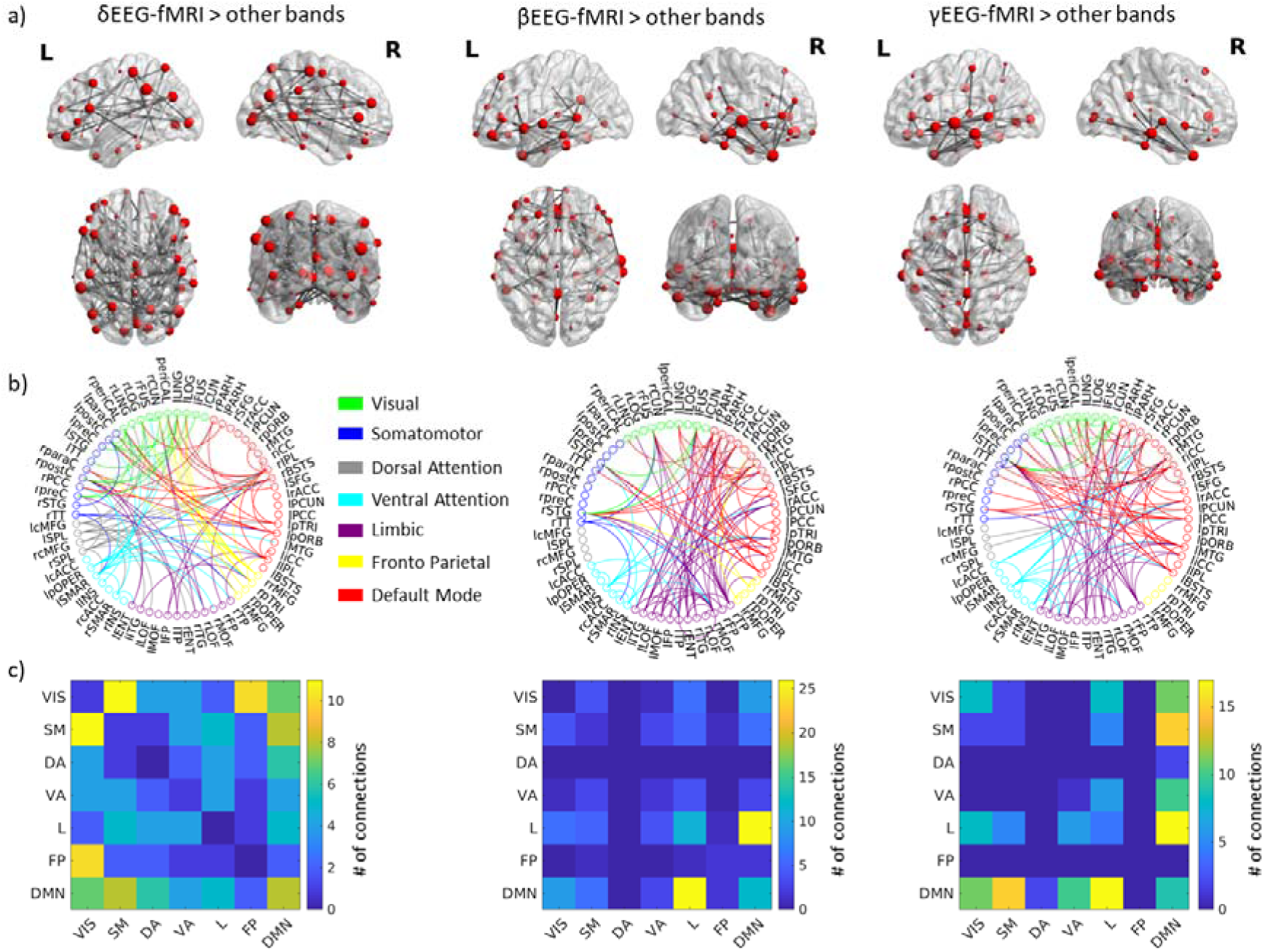
Frequency-specificity of temporally linked EEG-fMRI connectivity dynamics. a) Topographical distribution of the top-100 connections in which connectivity dynamics were more strongly linked between fMRI and δEEG than other EEG bands (left), between fMRI and βEEG than other EEG bands (center), and between fMRI and γEEG compared to other bands (right) (determined by NBS, 5000 iterations, cf. table 1). Color code and region labels correspond to Fig. 4. a) Location (center of gravity) of brain regions comprised in the respective top-100 connections. Sphere size depicts the number of the region’s connections among the top-100 cluster. b) The count of connections among the top-100 falling between a given pair of canonical intrinsic networks. The δ-dominated set most dominantly comprises connections of the Visual network to Somatomotor and Frontoparietal networks, as well as connections of the DMN especially to itself and Somatomotor areas. The β-dominated set of connections has a largely ventral distribution, most prominently comprising connections between the DMN to the Limbic network. The γ-dominated set predominantly contains connections between the DMN to the Somatomotor and Limbic networks. Results are visualized for the primary dataset, while frequency-specificity for δ, β and γ bands was likewise observed in the generalization dataset (table 1, SI Figure 3).

## Discussion

Temporal dynamics of neural communication is necessary for human behavior, since behavior is inherently flexible. While connectivity dynamics are most commonly studied using fMRI, the relationship between fMRI-derived whole-brain connectome reconfigurations and the dynamics of fast neurophysiological connectivity is still unclear. In this simultaneous EEG-fMRI study, we compared fMRI- and EEG-derived dynamic functional connectivity (dFC) concurrently across all cortical region pairs of the connectome. We demonstrated that the timecourse of fMRI-derived dFC shares mutual information with the timecourse of EEG-derived dFC across a vast proportion of all cortical region pairs. Interestingly, the cross-modal relationship of dFC was not dominated by any particular EEG frequency band, although the relationship progressively weakened with increasing EEG frequency when data points were limited (generalization dataset). In spite of the extensive spatial distribution of the cross-modal relationship in all frequencies, the region-pairs with the strongest tie between EEG- and fMRI-based connectivity dynamics varied across oscillation-bands (frequency-specificity in 21.8% of region pairs of the main dataset and 14.9% of the generalization dataset). These results provide strong evidence that fMRI-derived dFC patterns are directly linked to neurophysiological dFC between corresponding region pairs distributed across the whole-brain connectome.

### Static connectivity

In line with our prior work, the spatial pattern of static EEG and fMRI connectomes were significantly correlated (Sl Fig. 1). Only three prior studies have investigated whole-brain connectomes in concurrent EEG and fMRI (6, 11, 36), and the current study confirms this important observation. Interestingly, as in the prior studies, the spatial correspondence of the static connectivity architecture across fMRI and EEG was not restricted to any single oscillation frequency; rather, it was spatially extensive in all EEG frequency bands. However, the three prior studies were limited to static relationships, while the core advantage of concurrent EEG-fMRI recordings lies in its potential to reveal dynamics co-fluctuating across modalities.

### Connectivity dynamics of neural origin in both EEG and fMRI

Dynamic FC has been shown to exist both on slow hemodynamic (14) and fast electrophysiological timescales (47, 48). Regarding the relationship between EEG and fMRI, Chang et al. (27), Allen et al. (30) and Lamoš et al. (29) showed that BOLD FC dynamics are linked to band-limited EEG power in all or subsets of electrodes. The current study advances beyond EEG power to source-localized EEG connectivity, permitting edge-wise cross-modal comparison of connectivity dynamics over the whole connectome. This approach revealed that oscillatory coherence between brain regions in canonical EEG frequencies, i.e. fast electrophysiological connectivity, exhibits time-varying changes in the temporal range of infraslow BOLD connectivity dynamics. Our core finding is that the slow changes in EEG-derived FC temporally co-occur with changes in fMRI-derived FC across vast proportions of region pairs. The strength of this cross-modal correspondence varied over connections by EEG frequency (see section on frequency-specificity below).

This finding has implications for the interpretation of fMRI-derived dFC. Since physiological and non-physiological noise contribute to fMRI-derived dFC (49), the degree to which such dynamics reflect changes in neural communication is difficult to assess. Further, choosing an appropriate null model can be challenging (15, 23). Instead of comparing fMRI dFC to a null model, we chose to compare fMRI dynamics to a direct measure of neural dynamics, i.e. EEG. Our approach of concurrently assessing dFC in EEG provides evidence that veridic dynamics can indeed be derived from fMRI, as they are directly related to slow changes in the underlying electrophysiological connectivity. Another important advantage of our approach is the independence of outcomes from both EEG- and fMRI-related artefacts (e.g. eye movements in EEG or magnetic field inhomogeneities in fMRI), since spurious dynamics due to random noise in each modality will cancel each other out in the joint model we utilized. Comparing a spatially unspecific EEG parameter such as band-limited sensor-level power to local fMRI connections (27, 29, 30) is an approach prone to spurious cross-modal links stemming from a connection-unspecific global shift (e.g. breathing (50)). In contrast, our approach comparing both modalities on a connection-wise level is unlikely to be impacted by averaged global patterns.

Our results also have important implications for neurophysiological connectivity dynamics with respect to methodological reliability and observed timescales. Due to concerns regarding the ill-posed nature of EEG source localization, connectivity approaches in whole-brain parcellation space are underused (35). The close connection-wise relationship to fMRI-derived dFC provides strong support for the relevance of source-localized EEG to the study of the whole-brain functional connectome. With respect to timescales, prior MEG-based whole-brain investigations have established fast dynamics in interregional connectivity at ~50-100ms (47, 48). Connectivity dynamics at these fast timescales in EEG have been shown to correlate with the slow changes observed in fMRI, albeit in EEG sensor space rather that reconstructed brain parcellations (51, 52). Concurrent EEG-fMRI studies investigating neurophysiological dynamics at infraslow speeds have been either limited to a coarse global field average of connectivity across EEG electrode pairs (24, 25), or have focused solely on amplitude fluctuations (12). Extending beyond these important studies, we show that connectivity derived from fast oscillation phase coupling in EEG exhibits meaningful fluctuations in the infraslow range.

### Spatial distribution of the cross-modal dynamics

Interestingly, connections with the strongest interrelation between EEG and fMRI connectivity predominantly spanned across the canonical intrinsic networks. While we found that the vast majority of connections were dynamically linked between EEG and fMRI, the cross-modal relation was weaker within as compared to across networks with the exception of DMN-DMN connectivity. This finding may seem counterintuitive since the investigation of EEG amplitude fluctuations (as opposed to connectivity) shows a strong link between EEG and fMRI within ICNs (53, 54). However, while amplitude fluctuations may be suggestive of strong static within-network connectivity, they don’t inform about connectivity dynamics. Studying dynamics of amplitude correlations in MEG, De Pasquale et al. (55) observed strong connectivity dynamics both within and between networks for α- and β-bands. One likely explanation for our results is that within-network connections are less dynamic than connections between different intrinsic networks.

Indeed, this interpretation is in line with the observation that DMN regions are among the most dynamic, as measured by fMRI-derived dFC (15). We found the strongest relationship between EEG and fMRI dynamics in cortico-DMN as well as Somatomotor-Visual connections. The dominance of DMN interactions with other ICNs has been previously demonstrated for dynamic connectivity derived from MEG (55). Similarly, Vidaurre et al. (48) observed increased MEG coherence in a higher-order network comprising both DMN and Somatomotor-Visual connections. More generally, the DMN has been proposed to form a central hub that integrates multisensory input from different brain regions (56). This aligns with our observation that connections between DMN and the Visual and Somatomotor systems showed some of the strongest relations between EEG and fMRI dFC.

### Frequency-specificity of the cross-modal dynamics

We found both convergence and divergence between the different EEG frequencies; we observed a spatially widespread significant link to fMRI dFC for all EEG bands (Fig. 2), but also a considerable difference in the strength of this cross-modal relationship between bands across connectome space. Specifically, when directly contrasting frequency bands, we observed that the link between EEG and fMRI dFC was particularly strong for the δ band compared to all other bands in Somatomotor-Visual connections, and for β and γ bands in DMN-Limbic connections (Fig. 5). In the following, we discuss how the similarity of our findings across frequencies as well as the frequency-specific differences fit with prior literature.

The relationship between static MEG/EEG and fMRI connectivity mirrors our observations in the dynamic framework, namely that the cross-modal relationship is at the same time spatially widespread in all frequencies and exhibits variations across connections depending on neurophysiological frequency (8). Brookes et al. (5) showed that static intra-network connectivity of ICNs (no whole-brain approach) best correlates between MEG and fMRI for α and β bands (but no frequency showed a link between fMRI and MEG for the Visual network). In accordance with this study, we observed that EEG-fMRI co-dynamics within the DMN (the only ICN with strong cross-modal link for intra-network connections) are most prominent in the β band (Figure 4). However, contrasting Brookes et al. (5), Hipp and Siegel (8) stress that after taking into account the different SNR levels of the different frequency bands, the static whole-brain connectome is linked across MEG and fMRI over a broad frequency range from 2-128 Hz. A significant correlation between static connectivity in fMRI and all EEG bands is also backed by the results of Deligianni et al. (6) and our prior study (11) showing spatially distributed correlation of static connectivity across fMRI and all canonical EEG bands (replicated in the current study in both datasets; Sl Fig. 1). Importantly, in spite of the widespread cross-modal correspondence of static connectivity in all frequencies, Hipp and Siegel (8) report that the strength of this correspondence varies over frequencies for 21% of all connections. This ratio is very close to the 21.8% connections for which we report significant frequency-specificity in the dynamic framework.

On the dynamic timescale, less work compares frequency-specific patterns of the cross-modal relationship. At first glance, previous findings mapping band-limited EEG power (not connectivity) to fMRI dFC seem to promote a spatially localized and frequency-confined cross-modal relationship, but a closer look paints a picture of a spatially widespread relationship comprising all EEG frequencies. FMRI dFC correlates of α power, the most extensively studied band in this context, is widespread over several ICNs. Scheeringa et al. (57) showed that dynamic posterior α power co-fluctuates with fMRI-derived dFC within the Visual system, but also spreads to frontal and temporo-parietal regions mostly comprising the DMN. Chang et al. (27) showed that the link between EEG α power dynamics to fMRI-derived dFC extends beyond the Visual system to the DMN and Dorsal Attention ICNs. Tagliazucchi et al. (28) observed anticorrelation of central α to fMRI dFC of central regions. Extending this existing work to EEG connectivity dynamics, we demonstrate (in two independent datasets) that the link between fMRI-derived and α dFC spreads to the whole brain albeit with strongest effect size in Visual-Somatomotor connections. Chang et al. (27) extended investigations of EEG power beyond the α band to the θ frequency and showed that the variance of θ power is linked to dFC in all the ICNs they investigated (DMN, Dorsal Attention and Somatomotor). Finally, investigating all frequency bands, Tagliazucchi et al. (28) observed correlation of fMRI-derived dFC and fronto-central γ power dynamics, and correlation of fMRI-derived dFC with the centrally located dynamic α and β power. Though Allen et al. (30) found strong cross-modal effects for α power (that also comprised high θ), they further observed an occipitally centered fMRI dFC state correlated to central δ/θ EEG power, which interestingly was also the most commonly occurring state. Lamoš et al. (29) observed that θ power is linked to fMRI-derived dFC in the Visual ICN, whereas α and β power were linked to Somatomotor and attentional ICNs. Exploring intracranial EEG-fMRI dynamics, Ridley et al. (33) observed correlated variance of dFC in all frequency bands especially at higher frequencies such as γ. Taken together, although studies of EEG power cannot be directly compared to our study of EEG connectivity dynamics, the wide frequency range of the reviewed findings in that literature is in line with our observations regarding co-dynamics of fMRI and EEG connectivity

Further, this conclusion is backed by animal recordings finding electrophysiological correlates of dFC across all frequencies (albeit with very limited spatial coverage (31, 32)). The wide frequency range becomes most apparent in a comprehensive review of the respective literature (58). In conclusion our study not only confirms findings of previous studies with respect to the wide range of frequencies, but also extends them to the whole-brain scale, and most importantly to a connection-wise comparison of fMRI dFC to EEG connectivity dynamics. This study demonstrates for the first time that fMRI dynamics co-fluctuate with electrophysiological connectivity dynamics across all frequencies, and that those connectivity co-fluctuations are not driven by the dominant power of the α band. Indeed, we find more stable effects in terms of generalizability between the two datasets in lower frequencies (δ and θ, cf. Fig. 3) in line with previous work (27, 30).

### Methodological considerations and limitations

The relatively low spatial resolution (number of regions) of the selected Desikan atlas was imposed by having no more than 64 EEG electrodes. While it has been shown that a parcellation adapted to the actual EEG montage improves the quality of the source reconstruction (59), it is unclear if the fMRI signal would suffer from a parcellation scheme imposed by the EEG montage. A future approach could be extending the MR-compatible EEG setup to 128 or 256 electrodes (60) to gain data quality comparable to MEG recordings (61).

Regarding temporal resolution, the sliding window approach has been previously criticized for assuming slowly changing dynamics as opposed to instantaneous switches (see (62) for review). While the timescale of observable dynamics depends on the choice of window length, our 1-min window was chosen based on careful consideration of published recommendations (22, 63) to strike a balance between maximizing the number of datapoints (reliability of FC) in each window, and keeping the window short enough to detect FC dynamics at relevant timescales (see Methods). Future multi-modal studies with accelerated fMRI sequences (particularly simultaneous multislice acquisitions) will allow for a comparative analysis of different windowing parameters. Importantly, we show that results from the chosen parameter set generalize across two independent datasets, speaking to the robustness of the findings with the current window size.

Due to the more limited sample size and recording duration of the generalization dataset, β-and γ-connectivity seem to be influenced by the low signal-to-noise ratio of the generalization dataset (with 84% and 87% of βEEG- and γEEG-derived connections sharing no significant dynamics with fMRI, as opposed to virtually no insignificant connections in the primary dataset). Beyond the signal-to-noise limitations in human concurrent EEG-fMRI, there is evidence that γ-and β-band connectivity may contain information complementary to fMRI-derived connectivity. A weaker relation to fMRI connectivity has been reported for β and low-γ than for other bands in intracranial electrophysiological recordings in humans (33, 64) and animals (32, 65). The weaker relationship for β-and low γ-bands likely reflects a general property of the electrophysiology-fMRI relationship. This interpretation is also in line with our previous finding that γ-connectivity shows a unique relationship to structural connectivity not shared by fMRI-derived connectivity (11). Importantly, at the coarser ICN-wise (Fig. 4) as opposed to connection-wise resolution (Fig. 3), the results generalized across all bands including β and γ. The generalization of effects in this study is especially supportive of the robustness of the EEG-fMRI dFC relationship in light of substantial differences across the two datasets (3T vs. 1.5T MRI field strength, eyes-closed vs. eyes-open resting-state and differences in fMRI sequences and subject demographics).

### Conclusion

We observed a link between electrophysiological and fMRI-derived dFC across a large proportion of the connectome’s region pairs. This observation demonstrates that fMRI-derived connectivity captures infraslow dynamics of fast electrophysiological phase coupling. While the cross-modal link of dFC exists across all canonical electrophysiological frequency bands in a spatially distributed fashion, the strength of the cross-modal relationship varies over connections in a frequency-specific manner. The cross-modal tie was strongest in Visual to Somatomotor connections for the slower EEG-bands, and in connections involving the DMN (especially to the Limbic network) for faster EEG-bands. In conclusion, this study provides strong multimodal evidence for the existence of infraslow time-varying intrinsic connectivity dynamics across the connectome. This finding supports the functional importance of intrinsic connectivity dynamics in neural information processing and motivates taking such whole-brain connectivity reconfigurations into account when studying the foundations of cognition or clinical symptoms, be it with fMRI or neurophysiological methods such as EEG and MEG.

## Methods

We analyzed a primary dataset (n=26) and tested for generalizability across a different sample by using an independent dataset (n=16) from a different site. The generalization dataset is openly available at https://osf.io/94c5t/ and described in detail in Deligianni et al. (6, 36).

### Primary Dataset

#### Subjects

We recruited 26 healthy subjects (8 females, mean age 24.39, age range 18-31) with no history of neurological or psychiatric illness. Ethical approval has been obtained from the local Research Ethics Committee (CPP Ile de France III) and informed consent has been obtained from all subjects.

#### Data Acquisition

We acquired three runs of 10 minutes eyes-closed resting-state in one concurrent EEG-fMRI session (Tim-Trio 3T, Siemens). FMRI parameters comprised 40 slices, TR=2.0s, 3.0×3.0×3.0mm, TE = 50ms, field of view 192, FA=78°. EEG was acquired using an MR-compatible amplifier (BrainAmp MR, sampling rate 5kHz), 62 electrodes (Easycap), referenced to FCz, 1 ECG electrode, and 1 EOG electrode. Scanner clock was time-locked with the amplifier clock (66). Additionally, an anatomical T1-weighted MPRAGE sequence was acquired (176 slices, 1.0×1.0×1.0 mm, field of view 256, TR=7min).

The acquisition was part of a study with two additional naturalistic film stimulus of 10 minutes not analyzed in the current study, and acquired after runs 1 and 2 of the resting state as described in Morillon et al. (67). The three runs resulted in a total length of 30 minutes of resting-state fMRI per subject. Subjects wore earplugs to attenuate scanner noise and were asked to stay awake, avoid movement and close their eyes during resting-state recordings. In three subjects, one of three rest sessions each was excluded due to insufficient EEG quality.

#### Data processing

##### Atlas

T1-weighted images were used to delineate 68 cortical regions of the Desikan atlas (38, 39) and to extract a gray matter mask (recon-all, Freesurfer suite v6.0.0, http://surfer.nmr.mgh.harvard.edu/).

##### fMRI

The BOLD timeseries were corrected for slice timing and spatially realigned using the SPM12 toolbox (revision 6906, http://www.fil.ion.ucl.ac.uk/spm/software/spm12). Mean white matter and cerebrospinal fluid timecourses were extracted from a manually defined spherical 5mm ROIs using MarsBaR (v0.44, http://marsbar.sourceforge.net/). Using the FSL toolbox (v5.0, https://fsl.fmrib.ox.ac.uk/fsl/) the scull-stripped T1 image (fsl-bet), Desikan atlas, and grey matter delineation were linearly coregistered into the subject space of the T2* images (fsl-flirt v6.0). The fMRI timeseries were averaged for each of the 68 atlas regions, and the six movement parameters (from realignment), signal of CSF and white matter ROIs and grey matter global signal were regressed out of the region-wise timeseries. The resulting timeseries were bandpass-filtered at 0.009-0.08 Hz (68).

##### EEG

EEG was corrected for the gradient artefact induced by the scanner using the template subtraction and adaptive noise cancelation followed by lowpass filtering at 75Hz, downsampling to 250Hz (69) and cardiobalistic artefact template subtraction (70) using EEGlab v.7 (http://sccn.ucsd.edu/eeglab) and the FMRIB plug-in (https://fsl.fmrib.ox.ac.uk/eeglab/fmribplugin/). Data was then analyzed with Brainstorm software (42), which is documented and freely available under the GNU general public license (http://neuroimage.usc.edu/brainstorm, version 10^th^ August 2017). Bandpass-filtering was carried out at 0.3-70 Hz. Data was segmented according to TR of the fMRI acquisition (2s epochs). Epochs containing head motion artefacts in EEG were visually identified after semi-automatically preselecting epochs where signal in any channel exceeded the mean channel timecourse by 4 std. These segments were excluded from the analysis. Electrode positions and T1 were coregistered by manually moving the electrode positions onto the electrode artefacts visible in the T1 image. Using the OpenMEEG BEM model, a forward model of the skull was calculated based on the individual T1 image of each subject (41, 71).

The EEG signal was re-referenced to the global average and was projected into source space using the Tikhonov-regularized minimum norm (40) with the Tikhonov parameter set to 10% (Brainstorm 2016 implementation, assumed SNR ratio 3.0, using current density maps, constrained sources normal to cortex, depth weighting 0.5/max amount 10). Source activity was averaged to the regions of the Desikan atlas. For each epoch (length 2s) imaginary coherence of the source activity was calculated between each region pair (70) at 2Hz frequency resolution. The 2Hz bins were averaged for 5 canonical frequency bands: δ (0.5-4Hz), θ (4-8Hz), α (8-12Hz), β (12-30Hz), γ (30-60Hz).

##### Joint motion scrubbing

For all analyses, both fMRI volumes and EEG epochs were excluded for time periods where motion was identified in either modality. Time periods with motion were defined as volumes exceeding the framewise displacement threshold FD=0.5 in fMRI (72), and by visual inspection in EEG as described above. Additionally, for sliding window connectivity (see section Sliding window connectivity below), windows with more than 10% of their datapoints (>3 fMRI volumes or >3 EEG epochs) removed by this motion scrubbing procedure were excluded from dynamic connectivity analysis. The joint motion scrubbing approach resulted in a mean of 544 out of 870 sliding windows (range 262-813) for the main dataset and 216 out of 272 sliding windows (range 112-259) for the generalization dataset.

#### Connectivity

##### Static connectivity

Static connectivity was estimated for fMRI data by calculating Pearson’s correlation of the BOLD timecourse between each region pair over the duration of each run and averaged across the 3 runs. For EEG, the connectivity (imaginary coherence) calculated for each 2s epoch was averaged across all runs (Fig. 1a).

##### Sliding window connectivity

dFC matrices were calculated using a rectangular sliding window of 1 min. We averaged imaginary coherence of all 2s epochs over the 60s window for EEG (average over 30 datapoints per window), and used Pearson’s correlation over the 60s timecourse for fMRI (correlation over 30 datapoints at a TR of 2s per window). The window length was chosen to capture infraslow dynamics characteristic of fMRI dFC within limits of the following methodological considerations. The window size represents a tradeoff between maximizing the number of datapoints without discarding relevant dynamic BOLD frequencies (68, 73, 74) while also taking into account the theoretical limitations of shorter window lengths to reliably detect dFC (22, 63).

Leonardi and van de Ville (22) propose a rule of thumb choosing a window length exceeding the longest wavelength of the BOLD signal around 100s. This rule of thumb has been confirmed by Zalesky et al. (63), but the authors also noted that a dynamic signal can still be detected (with lower power) for shorter window lengths (40s such as used in Shirer et al. (74)). Shine et al. (73) showed that those theoretical limits can be undersampled down to 10s (14 data points in their data). Chang et al. (27) find a significant relationship between fMRI dFC and global α power dynamics peaking at 30s to 55s, and θ power dynamics peaking at window sizes between 65s and 70s. Given these results from prior studies we consider the 60s window (30 data points) a good tradeoff between being able to detect dynamics and having high detection power.

Most importantly, we show that our results reliably replicated across two independent datasets with the chosen parameter set.

Normalized mutual information defined by 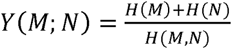 with H(M), H(N) being the entropies of observations N and M and H(M,N) the joint entropy (75, 76) was calculated for the resulting EEG and fMRI dFC matrices, building a new connectivity matrix of joint EEG-fMRI, based on mutual information strength between the modalities (for each EEG frequency band). In contrast to linear measures such as correlation, mutual information is an information theoretic measure which is able to also capture cross-modal relationships in connectivity dynamics without assuming linearity or Gaussian constraints (77). Mutual information has previously been shown to be helpful when combining EEG and fMRI (78).

##### Null model

For each connection, mutual information was then compared to a null model of EEG-fMRI mutual information (Fig. 1b) using temporally phase-randomized fMRI dFC timecourses of each connection (45, 46). Specifically, we applied the approach of phase-randomization proposed by Theiler et al. (45) to whole-brain connectomes by Fourier transforming the fMRI dFC timecourse at each connection. Then the phases of this transformation were randomly shifted. The result of the phase-shift was then back-transformed using the inverse Fourier transform. Importantly, this operation will create surrogate data with Fourier spectra and autocorrelation equal to that of the original timecourse. Note that unlike unimodal studies of fMRI dFC that are interested in the relationship between two fMRI signal timecourses and thus phase randomize the signal timecourse (e.g. (23)), we are interested in the relationship across EEG and fMRI connectivity timecourses, hence randomizing the connectivity timecourse for one modality.

The randomization process was carried out 50 times for the fMRI dFC timecourse for each subject and connection, and mutual information between the phase-randomized fMRI dFC timecourse and the unaltered EEG dFC timecourse was calculated for each iteration. For group-level statistical comparison of this null model to the original EEG-fMRI mutual information, we calculated an average null mutual information matrix from the 50 iterations for each subject. We subjected the original mutual information matrix to connection-wise paired t-tests against the null model over subjects. The ensuing p-values were Bonferroni corrected for the number of connections. This procedure was repeated for each EEG frequency band.

##### Intrinsic network analysis

To further interpret outcomes in the context of neurocognitive networks, we mapped the extracted 200 connections to 7 canonical ICNs (Visual, Somato-Motor, Default Mode, Fronto-Parietal, Dorsal Attention, Limbic, and Ventral Attention [largely corresponding to Cingulo-Opercular (80)]) as described in Yeo et al. (3). See Table S9 for the exact mappings between each brain region and ICN. The number of connections falling into each network pair were counted (e.g. DMN to Visual). To assure that the observed connectivity pattern did not arise from random sampling into the different networks, we also created 100,000 random networks of 200 connections to derive the probability that a connection randomly falls into one of the ICN pairs.

##### Frequency-specific analysis

To test for frequency-specificity, we included frequency-specific Fisher z-transformed mutual information matrices of all bands (δEEG-fMRI, θEEG-fMRI, etc.) and for all subjects in an ANOVA (frequency band as one factor with 5 levels). Note that this analysis compares EEG-fMRI mutual information matrices from different bands to each other, and as such does not require the above-described null model generated for the first statistical analysis. An F-Test was carried out at each connection to determine if the EEG-bands contributed differentially to the mutual information with fMRI-derived dFC. We used Network Based Statistics (NBS) to correct for multiple comparisons (79) (https://sites.google.com/site/bctnet/comparison/nbs, Version 1.2). NBS controls the family-wise error rate of the mass-univariate testing at every connection. This method is a non-parametric cluster-based approach to finding connected sets of nodes that significantly differ across thresholded connectivity matrices. Posthoc t-tests between one band vs. mean of all the other bands were carried out at each connection followed by NBS to explore if any EEG band expressed a network of stronger mutual information than observed in the other bands.

### Generalization dataset

#### Subjects

This dataset comprises 17 healthy adults. Ethical approval has been obtained from the UCL Research Ethics Committee (project ID:4290/001) and informed consent has been obtained from all subjects. One subject was excluded as T1 data quality was not sufficient to run the Freesurfer recon-all command, resulting in a final group of 16 subjects (6 females, mean age: 32.41, range 22-53).

#### Data Acquisition

We used one session of 10 minutes 48 seconds eyes-open resting-state (Avanto 1.5T, Siemens, 30 slices, TR=2.16s, slice thickness 3mm + 1mm gap, effective voxels size 3.3×3.3×4.0mm, TE = 30ms, field of view 210, flip angle 75 degrees) concurrent EEG-fMRI (63 scalp electrodes BrainCap MR, referenced to FCz, 1 ECG electrode). Scanner clock was timelocked with the MR-compatible amplifier (BrainAmp MR, sampling rate 1kHz) clock. A T1-weighted structural image was also obtained (176 slices, 1.0×1.0×1.0 mm, field of view 256, TR=11min). During the resting-state run, the subjects had their eyes open and were asked to remain awake and fixate on a white cross presented on a black background. Their head was immobilized using a vacuum cushion during scanning.

#### Data processing

The fMRI data was processed as described for the primary dataset with the exception that no slice-time correction was carried out (in accordance with the original processing in Deligianni (6)). EEG was corrected for the gradient artefact using the template subtraction and adaptive noise cancelation followed by a downsampling to 250Hz and cardiobalistic artefact template subtraction using the Brain Vision Analyzer 2 software (Brain Products, Gilching, Germany). Due to apparent low frequency drift artefacts in several subjects, EEG data was high pass filtered at 0.05Hz instead of the 0.03Hz used in the primary dataset. Because of the differing TR the sliding window for 1 minute was now consisting of 28 volumes (28*2.16s = 60.48s). EEG data processing was equivalent to the primary dataset, with the epochs being 2.16s instead of 2s to match the fMRI TR.

All following analysis steps were identical to the primary dataset.

## Acknowledgements

We thank Katia Lehongre and Benjamin Morillon (primary dataset) and Fani Deligianni and Jonathan Clayden (generalization dataset) for generously sharing their data. JW and SS were supported by a Beckman Institute MoCC seed grant. ALG was supported by ERC 260347 – COMPUSLANG.

## Data availability

Primary data will be made available by request to ALG. The generalization dataset is publicly available at https://osf.io/94c5t/. Custom analysis code is publicly available at https://github.com/jwirsich/dFC-EEG-fMRI.

## Author contributions

JW and SS designed the study, developed the methods and wrote the manuscript. ALG collected data.

## Supplementary Information

### SI: Results

#### Static connectivity

**SI Fig. 1:**
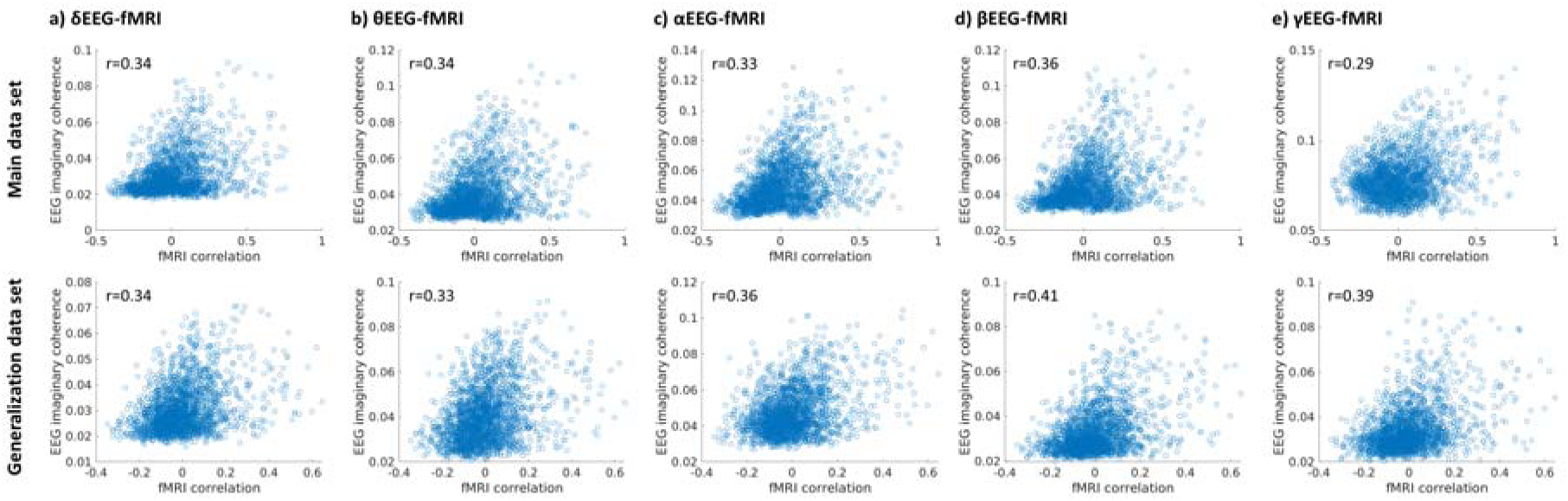
Correlation of EEG-fMRI static connectivities (averaged over each session and all subjects)

#### Mutual information strength

**SI Fig. 2:**
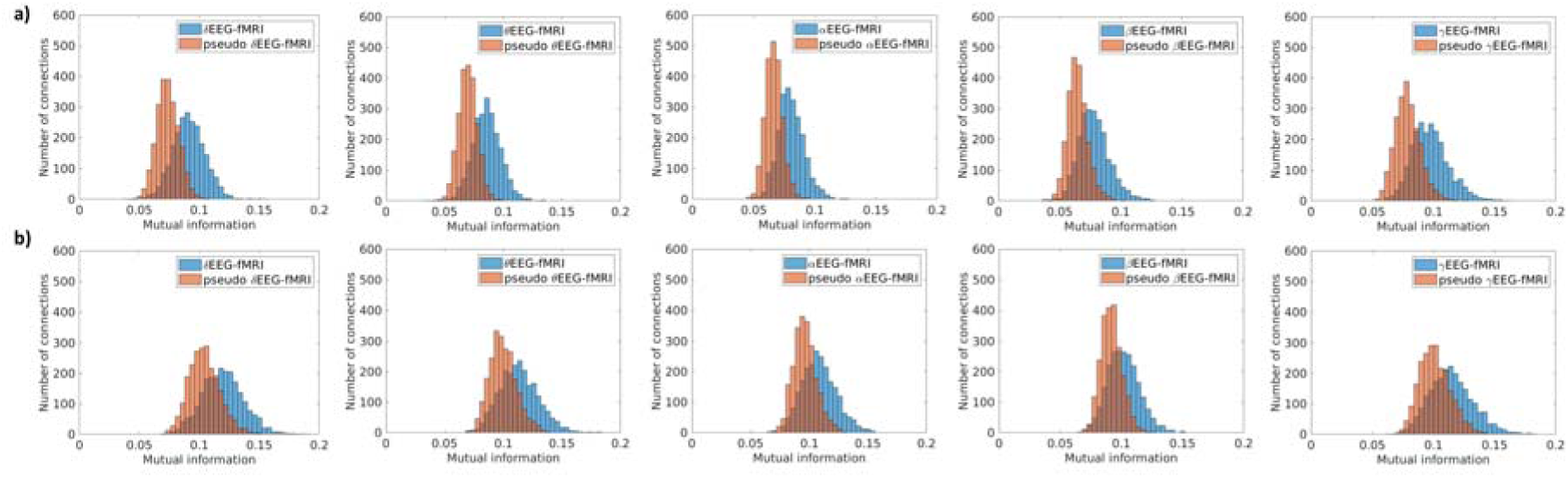
Distribution of mutual information across all connections between fMRI and EEG dFC time courses using real vs. phase-scrambled fMRI connectivity timecourses (null model) for each EEG band. Data are shown for a randomly selected subject of a) the main dataset and b) the generalization dataset. Comparison to the null distribution revealed that EEG and fMRI dynamics are significantly linked over time irrespective of EEG oscillation band for virtually all connections of the main dataset. A similar effect is observed for a large proportion of connections in the generalization dataset (see main manuscript Fig. 2).

When assessing generalization at a connection-wise resolution, we observed correlation of connection-wise EEG-fMRI mutual information for δ, θ, and α bands but not for β and γEEG (Fig. 3). To assess whether this discrepancy was due to lower signal to noise ratio in the higher frequencies, we investigated whether replication within each dataset would show a similar pattern. We divided the primary dataset (respectively the generalization dataset) into two groups of 8 subjects each (taking only the first 16 subjects of the primary dataset). Indeed, we observed a within dataset correlation of fMRI vs. δ/θ/α connection-wise mutual information, whereas no correlation between fMRI vs. β/γ was observed (Table S1). This outcome confirms that β and γ bands had lower SNR, which likely contributed to the lack of correlation of connection-wise mutual information across the primary and generalization datasets.

**Table S1:** Within group correlation of mutual information when splitting up subjects into two groups of 8 (for main dataset the first 16 subjects were taken to create equal groups of 8)

| | δEEG-fMRI | θEEG-fMRI | αEEG-fMRI | βEEG-fMRI | γEEG-fMRI |
| --- | --- | --- | --- | --- | --- |
| Main dataset | 0.51 | 0.36 | 0.19 | 0.04 | 0.03 |
| Generalization Data | 0.26 | 0.30 | 0.22 | 0.04 | 0.002 |

#### Impact of Movement

We took the precaution to remove movement confounds affecting fMRI and EEG by excluding all EEG-segments and fMRI volumes that either had above threshold framewise displacement (1) or showed motion artefacts on the EEG timecourse. The imaginary coherence used to measure EEG connectivity provides additional cleaning as it discards spurious zero-lag connectivity from global artifacts, which could be caused by residuals of movement, cardiobalistic and gradient artefacts. Nevertheless, as oscillations recorded in scalp EEG might be particularly impacted by head movement (2), we tested how much the framewise displacement value, a measure of head motion derived from raw fMRI volumes (see Methods), is correlated over time to the global connectivity of each EEG band (imaginary coherence timecourse over the 2s segments, averaged across all connections). Only segments included in the main analyses, i.e. those with low movement, were considered. We found no consistent evidence for a (linear) relationship between the framewise displacement and global EEG connectivity. Specifically, correlations were significant only for a small subset of sessions (table S2), but importantly at negligible effect sizes; Mean correlation (average across sessions) between the two measures was consistently low (peaking at R=0.04669 for gamma of the main data set).

**Table S2:** Correlation between EEG imaginary coherence timecourse (averaged across all connections) and framewise displacement (1) of fMRI volumes as a measure of head motion.

| | Mean R (SD) | Number of sessions reaching $p<0.05$ (Bonferroni corrected by total no. of sessions) |
| --- | --- | --- |
| **Main data set** |  |  |
| δEEG | 0.0019 (0.07447) | 0/75 |
| θEEG | -0.0040 (0.07669) | 1/75 |
| αEEG | -0.02038 (0.06652) | 1/75 |
| βEEG | 0.03100 (0.09112) | 3/75 |
| γEEG | 0.04669 (0.1178) | 11/75 |
| **Replication data set** |  |  |
| δEEG | 0.02769 (0.08942) | 2/16 |
| θEEG | 0.02125 (0.06674) | 0/16 |
| αEEG | 0.02652 (0.06836) | 0/16 |
| βEEG | 0.03418 (0.06376) | 0/16 |
| γEEG | 0.02217 (0.07287) | 0/16 |

#### Non-random nature of mutual information distribution across canonical ICNs

We established that the distribution of the top-200 connections and the ensuing DM, VIS and SM network dominance (Fig. 4) were not driven by the number of ICN nodes or other potential biases. For each network pair (e.g. DMN-VIS) we tested whether the number of connections was significantly higher than chance by randomly selecting (n=100,000 iterations) 200 connections from the main dataset (Table S3-S7 for five EEG frequencies). Additionally, Table S8 represents an equivalent analysis for a global measure of each ICN’s connectivity to the entirety of the connectome by averaging all connections of the respective ICN irrespective of where they connect to.

#### Spatial correspondence of ICN-mapping across datasets

We observed that the topographic distribution of the top-200 connections (connections with strongest mutual information between EEG dFC and fMRI dFC) over intrinsic connectivity networks (ICNs, Fig. 4) was similar across datasets. We performed an additional permutation analysis (n=100,000 iterations) to ascertain that this similarity was not due to chance. Specifically, starting with the top-200 connections of both main and generalization datasets, we randomly reassigned which dataset each of these connections belonged to. Next, we calculated the number of connections across each ICN pair for the permuted data (equivalent to Fig. 4b). We then calculated the difference between ICN-ICN connection counts in the permuted versions of main and generalization datasets. Finally, for each ICN pair we compared this difference of counts to the difference calculated from the actual main and generalization datasets. The difference (dissimilarity) of connection counts was not statistically distinguishable between the original data and the permutation null distribution in any ICN pair for any band (p>0.0018 for all EEG frequency bands, Bonferroni corrected for multiple comparisons in 28 network pairs). This outcome indicates that the connection counts in the main and generalization datasets were drawn from the same distribution in all ICN pairs, showing that the strongest connections have comparable topographic distribution over ICNs in the two datasets.

**Table S3:** p-values of comparing the number of top-200 most significant EEG-fMRI connections for a given EEG band falling within an ICN-ICN pair as compared to randomly sampling 200 connections of the brain (100.000 iterations). Yellow cells show connections significant at p<0.0018 (p<0.05, Bonferroni corrected for 28 comparisons) in the main dataset. The green cells additionally replicated at p<0.05 uncorrected in the generalization dataset. p-values (as described in the above caption) for δEEG-fMRI

|  | VIS | SM | DA | VA | L | FP | DMN |
| --- | --- | --- | --- | --- | --- | --- | --- |
| VIS | 0.8992 | 0 | 0.0151 | 0.0005 | 0.6765 | 0.0044 | 0.7043 |
| SM |  | 0.4827 | 0.0247 | 0.1860 | 0.7815 | 0.5019 | 0.4624 |
| DA |  |  | 0.0095 | 0.1976 | 0.3664 | 0.0076 | 0.0291 |
| VA |  |  |  | 0.8407 | 0.8402 | 0.6865 | 0.2662 |
| L |  |  |  |  | 0.9787 | 0.9848 | 0.9999 |
| FP |  |  |  |  |  | 0.4067 | 0.0127 |
| DMN |  |  |  |  |  |  | 0.9635 |

**Table S4:** p-values of comparing the number of top-200 most significant EEG-fMRI connections for a given EEG band falling within an ICN-ICN pair as compared to randomly sampling 200 connections of the brain (100.000 iterations). Yellow cells show connections significant at p<0.0018 (p<0.05, Bonferroni corrected for 28 comparisons) in the main dataset. The green cells additionally replicated at p<0.05 uncorrected in the generalization dataset. p-values (as described in the above caption) for ϑEEG-fMRI.

|  | VIS | SM | DA | VA | L | FP | DMN |
| --- | --- | --- | --- | --- | --- | --- | --- |
| VIS | 0.9804 | 0 | 0.2328 | 0.0269 | 0.5434 | 0.2324 | 0.2944 |
| SM |  | 0.3024 | 0.0002 | 0.0093 | 0.9694 | 0.4980 | 0.8289 |
| DA |  |  | 0.0815 | 0.6904 | 0.7721 | 0.1429 | 0.0125 |
| VA |  |  |  | 0.8387 | 0.7109 | 0.1973 | 0.0365 |
| L |  |  |  |  | 0.9785 | 0.9179 | 0.9970 |
| FP |  |  |  |  |  | 0.4073 | 0.0626 |
| DMN |  |  |  |  |  |  | 0.9302 |

**Table S5:**
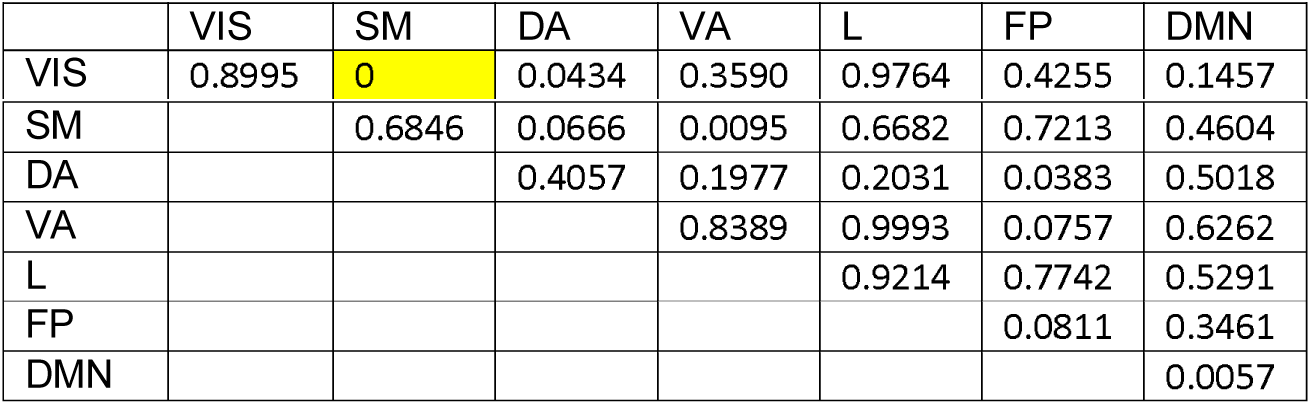
p-values of comparing the number of top-200 most significant EEG-fMRI connections for a given EEG band falling within an ICN-ICN pair as compared to randomly sampling 200 connections of the brain (100.000 iterations). Yellow cells show connections significant at p<0.0018 (p<0.05, Bonferroni corrected for 28 comparisons) in the main dataset. The green cells additionally replicated at p<0.05 uncorrected in the generalization dataset. p-values (as described in the above caption) for αEEG-fMRI

**Table S6:** p-values of comparing the number of top-200 most significant EEG-fMRI connections for a given EEG band falling within an ICN-ICN pair as compared to randomly sampling 200 connections of the brain (100.000 iterations). Yellow cells show connections significant at p<0.0018 (p<0.05, Bonferroni corrected for 28 comparisons) in the main dataset. The green cells additionally replicated at p<0.05 uncorrected in the generalization dataset. p-values (as described in the above caption) for ϐEEG-fMRI.

|  | VIS | SM | DA | VA | L | FP | DMN |
| --- | --- | --- | --- | --- | --- | --- | --- |
| VIS | 0.8990 | 0.1957 | 0.9698 | 0.3589 | 0.6776 | 0.9695 | 0.8639 |
| SM |  | 0.6820 | 0.5009 | 0.9987 | 0.2004 | 0.4996 | 0.3615 |
| DA |  |  | 0.4060 | 0.4104 | 0.7750 | 0.7507 | 0.8040 |
| VA |  |  |  | 0.2516 | 0.5525 | 0.4132 | 0.7398 |
| L |  |  |  |  | 0.1753 | 0.0018 | 0.0008 |
| FP |  |  |  |  |  | 0.0830 | 0.0128 |
| DMN |  |  |  |  |  |  | 0.0000 |

**Table S7:** p-values of comparing the number of top-200 most significant EEG-fMRI connections for a given EEG band falling within an ICN-ICN pair as compared to randomly sampling 200 connections of the brain (100.000 iterations). Yellow cells show connections significant at p<0.0018 (p<0.05, Bonferroni corrected for 28 comparisons) in the main dataset. The green cells additionally replicated at p<0.05 uncorrected in the generalization dataset. p-values (as described in the above caption) for γEEG-fMRI.

|  | VIS | SM | DA | VA | L | FP | DMN |
| --- | --- | --- | --- | --- | --- | --- | --- |
| VIS | 0.8974 | 0.0036 | 0.4256 | 0.1216 | 0.8847 | 0.8560 | 0.0951 |
| SM |  | 0.4841 | 0.0002 | 0.0495 | 0.4136 | 0.8923 | 0.0001 |
| DA |  |  | 0.4052 | 0.1989 | 0.9193 | 0.3886 | 0.2127 |
| VA |  |  |  | 0.8398 | 0.7127 | 0.4131 | 0.0372 |
| L |  |  |  |  | 0.8114 | 0.3653 | 0.7970 |
| FP |  |  |  |  |  | 0.4075 | 0.9676 |
| DMN |  |  |  |  |  |  | 0.9306 |

**Table S8:** p-values of comparing the number of top-200 most significant EEG-fMRI connections linked to a specific ICN as compared to randomly sampling 200 connections of the brain (100.000 iterations). In contrast to tables S3-S7, for each frequency all connections of a given ICN are aggregated irrespective of what other ICN they connect to. This provides a global measure of an ICN’s connectivity to the entirety of the connectome. significant connections are highlighted in green and yellow p<0.0071 (p<0.05, Bonferroni corrected for 7 comparisons). Green cells replicated at uncorrected threshold p<0.05 in generalization dataset.

|  | VIS | SM | DA | VA | L | FP | DMN |
| --- | --- | --- | --- | --- | --- | --- | --- |
| δEEG-fMRI | 0 | 0.0003 | 0.0001 | 0.1152 | 1 | 0.0569 | 0.9601 |
| θEEG-fMRI | 0 | 0 | 0.0040 | 0.0177 | 1 | 0.3252 | 0.7609 |
| αEEG-fMRI | 0.0262 | 0.0030 | 0.0360 | 0.6145 | 0.9994 | 0.4187 | 0.0671 |
| βEEG-fMRI | 0.9895 | 0.7436 | 0.9952 | 0.9696 | 0.0020 | 0.0571 | 0.0001 |
| γEEG-fMRI | 0.1685 | 0 | 0.0873 | 0.0598 | 0.9917 | 0.9947 | 0.0254 |

#### Frequency-specific networks and their mutual information strength

**SI Fig. 3:**
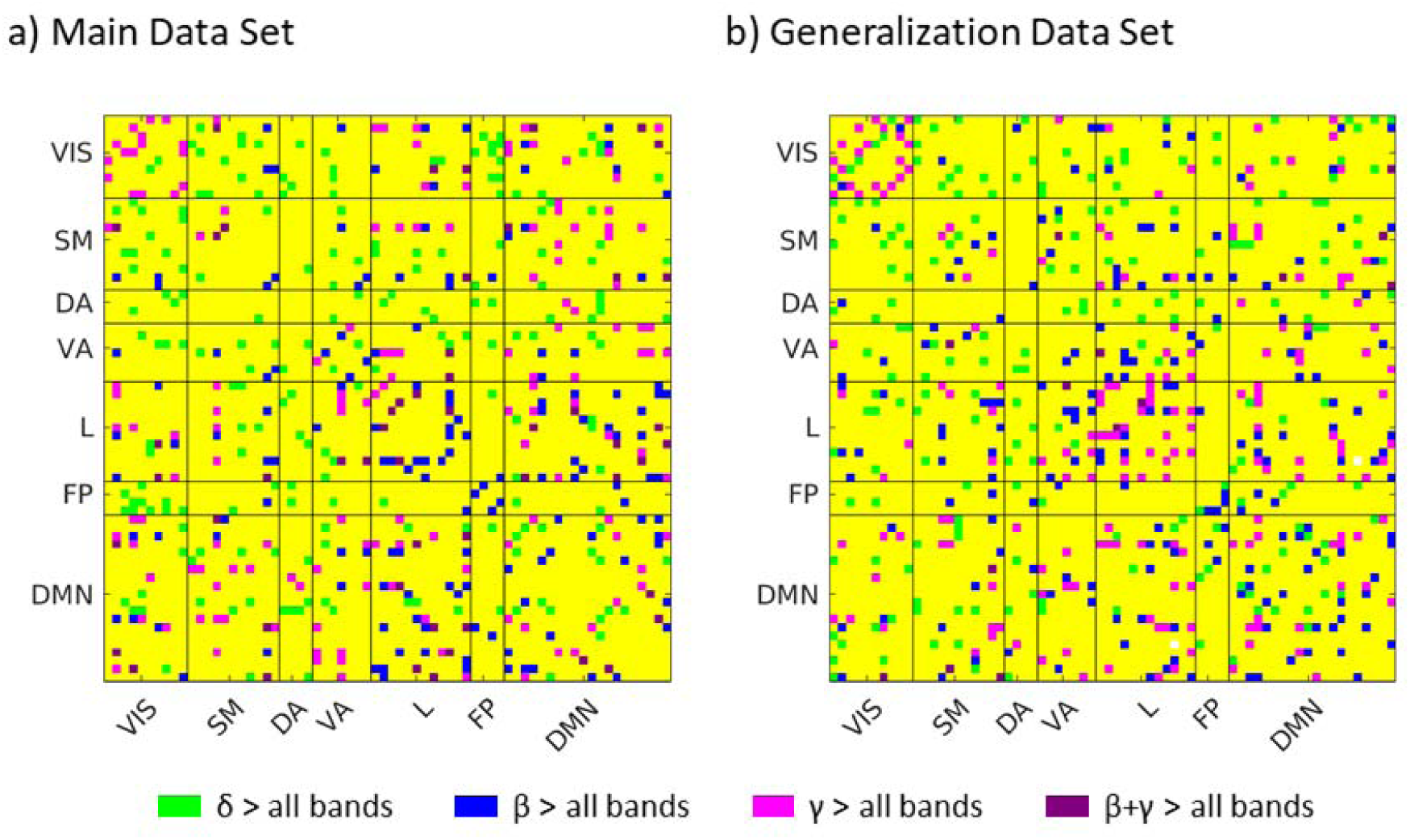
The outcome of post hoc t-tests of frequency-specificity (NBS-network limited to 100 connections, see Table 1). Plot a) depicts the connections shown in Fig. 5 but projected into one matrix for a direct spatial comparison. Plot b) shows equivalent data for the generalization dataset. Purple dots signify connections that were both significant for the ‘ϐ>all other bands’ and ‘γ>all other bands’ contrasts. No overlap was observed for δ with the other bands. Overall, the distribution for the δ, ϐ, and γ bands encompass largely different sets of connections.

To test if the mutual information values in the frequency-specific subnetworks (Fig. 5) are generally the highest values of the band-specific mutual information distribution, we determined whether for a given band mean mutual information was generally higher compared to the rest of the brain network. This is the case for all band-specific subnetworks (delta>other bands: p = 7.7*10^−41^, beta>other bands: p=8.0*10^−19^, gamma>other bands: p=2.8*10^−9^). This relationship also holds when masking the generalization dataset by the band-specific subnetworks of the main dataset (delta>bands: p=1.2*10^−5^, beta>bands: p = 2.0*10^−6^, gamma>bands: 0.0051), thus demonstrating the generalizability of the subnetworks selected as frequency-specific. This relationship also holds when masking the generalization dataset by the band-specific subnetworks of the main dataset (delta>bands: p=1.2*10^−5^, beta>bands: p = 2.0*10^−6^, gamma>bands: 0.0051), thus demonstrating the generalizability of the subnetworks selected as frequency-specific.

Further, when counting the number of the top-100 significant connections in each band-specific subnetwork over the seven canonical ICNs (cf. Fig. 5b), the distribution of the count (visualized in Fig. 5c) was correlated across datasets for delta, beta and gamma subnetworks: δEEG-fMRI/βEEG-fMRI/γEEG-fMRI: r=0.68/0.82/0.96, p=0.00062/4.7*10^−6^ /6.2*10^−12^. This outcome additionally supports spatial correspondence across datasets with respect to canonical neurocognitive networks for the three bands.

#### Atlas and ICN labels

**Table S9:**
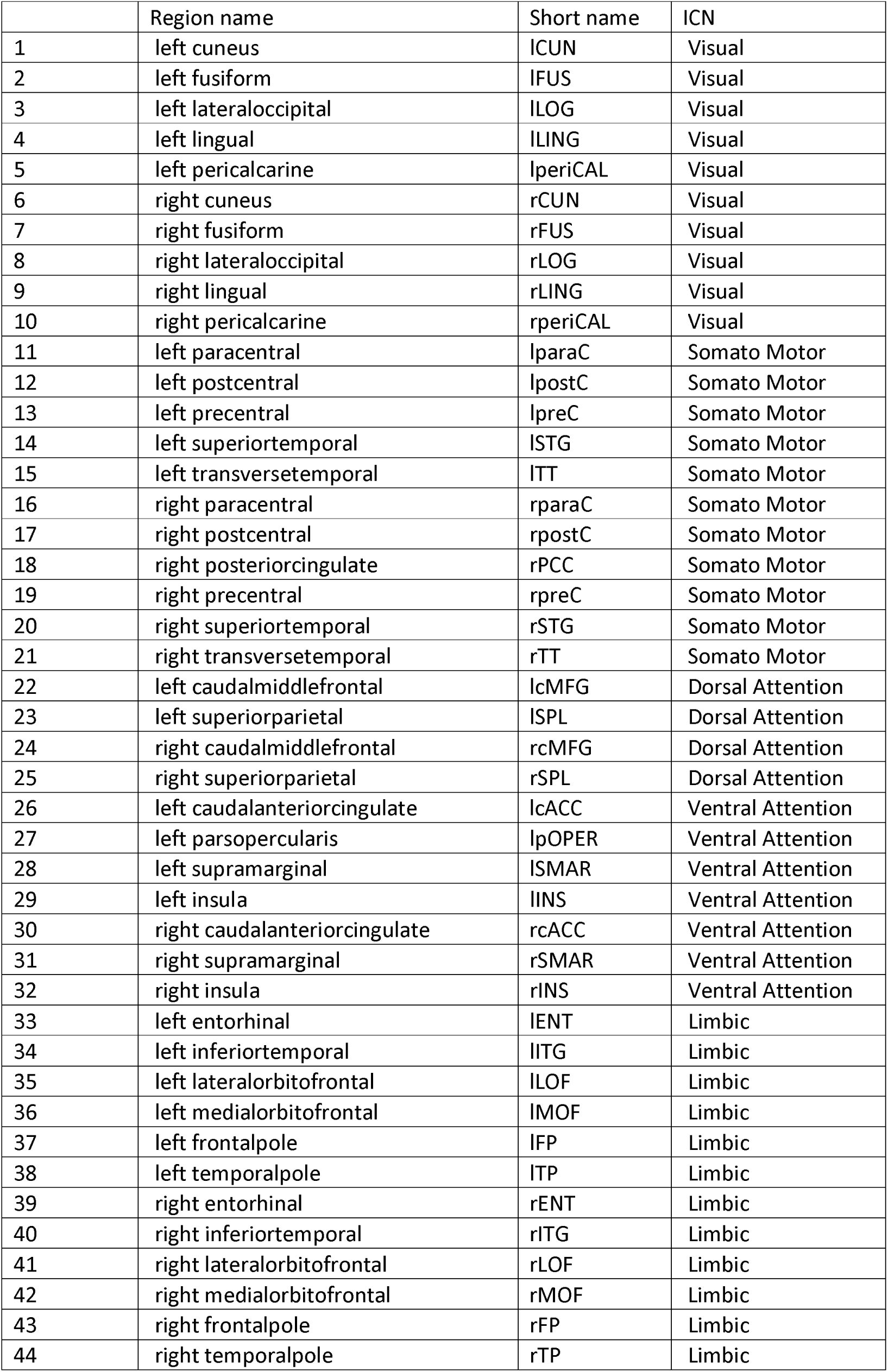
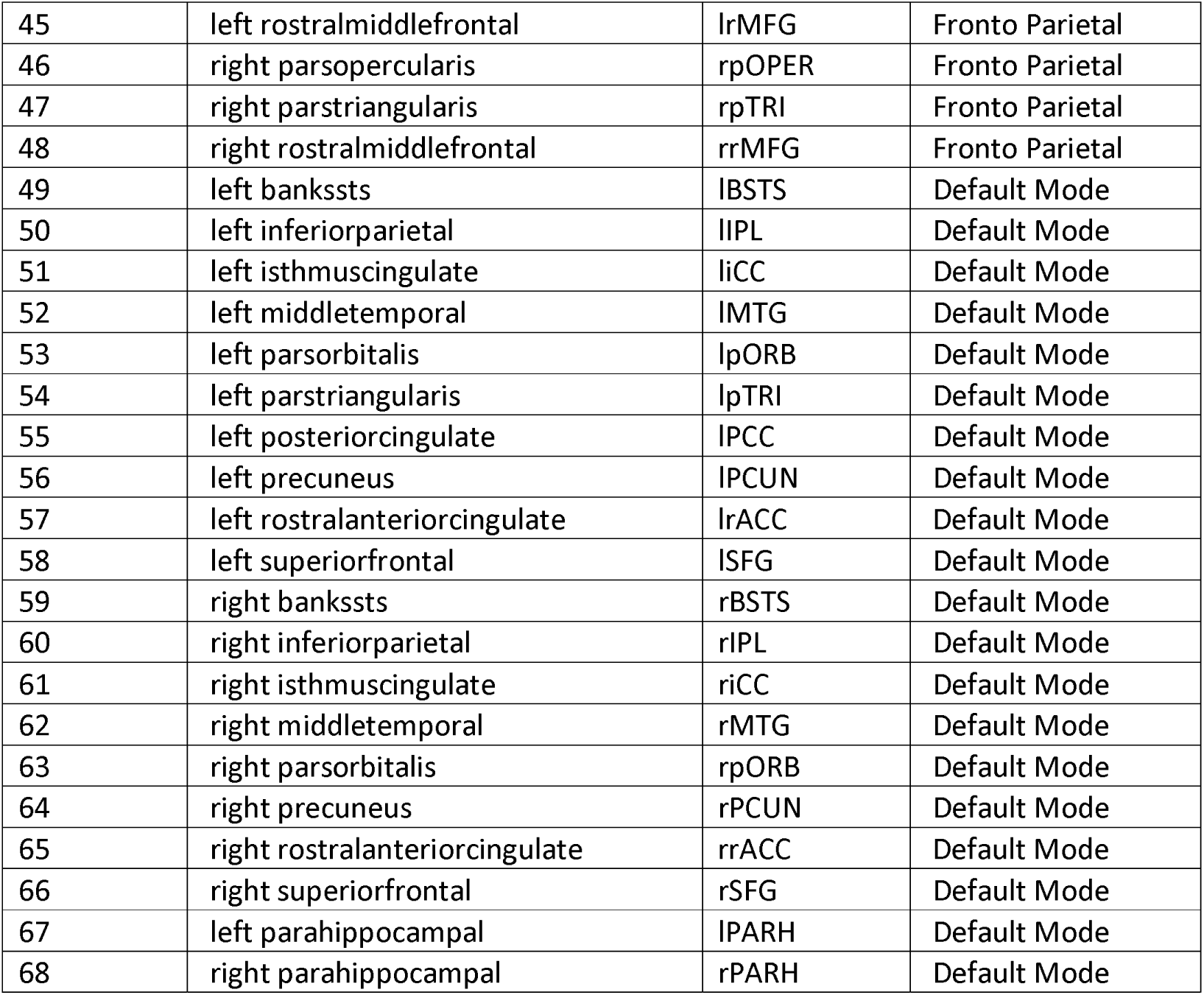
Regions of the Desikan atlas and their mapping to the canonical ICN networks

